# Validation of a new coil array tailored for dog functional magnetic resonance imaging (fMRI) studies

**DOI:** 10.1101/2022.06.14.496064

**Authors:** C.-N. Alexandrina Guran, Ronald Sladky, Sabrina Karl, Magdalena Boch, Elmar Laistler, Christian Windischberger, Ludwig Huber, Claus Lamm

## Abstract

Comparative neuroimaging allows for the identification of similarities and differences between species. It provides an important and promising avenue, to answer questions about the evolutionary origins of the brain’s organization, in terms of both structure and function. Dog fMRI has recently become one particularly promising and increasingly used approach to study brain function and coevolution. In dog neuroimaging, image acquisition has so far been mostly performed with coils originally developed for use in human MRI. Since such coils have been tailored to human anatomy, their sensitivity and data quality is likely not optimal for dog MRI. Therefore, we developed a multi-channel receive coil (K9 coil) tailored for high-resolution functional imaging in canines, optimized for dog cranial anatomy. In this paper we report structural (n = 9) as well as functional imaging data (resting-state, n = 6; simple visual paradigm, n = 9) collected with the K9 coil in comparison to reference data collected with a human knee coil. Our results show that the K9 coil significantly outperforms the human knee coil, improving the signal-to-noise ratio across the imaging modalities. We noted increases of roughly 45% signal-to-noise in the structural and functional domain. In terms of translation to functional fMRI data collected in a visual flickering checkerboard paradigm, group-level analyses show that the K9 coil performs better than the knee coil as well. These findings demonstrate how hardware improvements may be instrumental in driving data quality, and thus, quality of imaging results, for dog-human comparative neuroimaging.

**Significance Statement:** Comparative neuroimaging is a powerful avenue to discover evolutionary mechanisms at the brain level. However, data quality is a major constraint in non-human functional magnetic resonance imaging. We describe a novel canine head coil for magnetic resonance imaging, designed specifically for dog cranial anatomy. Data quality performance and improvements over previously used human knee coils are described quantitatively. In brief, the canine coil improved signal quality substantially across both structural and functional imaging domains, with strongest improvements noted on the cortical surface.

## 1. Introduction

Comparative neuroimaging aims to find the commonalities and differences in brains and brain function of different species. The focus of comparative neuroimaging often lies on great apes and other non-human primates (de Schotten et al., 2019; Rilling, 2014), but by focusing on comparisons between primates, insights on convergent evolution are limited. Convergent evolution describes the advent of a trait, such as a neural mechanism, in phylogenetically distant species, where both species developed the trait independently (e.g. wings in bats and birds). Neuroscience research and neuroimaging in birds (Behroozi, 2019; Behroozi et al., 2020; Güntürkün & Bugnyar, 2018) and reptiles (Behroozi et al., 2018) have shown that cognition is not reliant on the presence of a neocortex. Therefore, looking at sophisticated behaviors in more distant species outside the primate lineage should not be neglected and indeed non-primate neuroscience has seen a rise of interest in the past decades (Bunford et al., 2017; De Groof et al., 2013; Mars et al., 2016; Xu et al., 2020).

With regard to convergent evolution, dogs, C*anis lupus familiaris*, are a study species of the highest interest: they excel in social cognition, often outperforming great apes in their understanding of social cues from humans (Huber, 2016; Kaminski & Nitzschner, 2013; Kirchhofer et al., 2012). This places the dog at a prime position for investigating the evolution of social cognition and other cognitive skills, mirrored in an increase of neuroimaging studies of dogs in recent years (Berns, 2013; Bunford et al., 2017; Huber & Lamm, 2017; Thompkins et al. 2016, for reviews).

Dogs have the added advantage of being highly trainable, which makes it possible to perform awake, unrestrained and unsedated neuroimaging in dogs (Berns et al., 2012; Karl et al., 2019; Strassberg et al., 2019), opening the possibility for classical functional magnetic resonance imaging studies in this species, something that is not easily possible in rodents (e.g., Keilholz et al., 2004), birds, or monkeys without fixating, restraining, or sedating the animals.

However, many challenges for canine neuroimaging remain to be met. Training dogs to lie still and voluntarily stay in the scanner environment while being attentive to the presented stimuli is very time consuming (Berns et al., 2012; Karl et al., 2019; Strassberg et al., 2019). Canine neuroimaging runs also need to be shorter than those typically used in humans, and usually amount to a maximum length of 5 minutes, as even highly trained dogs cannot maintain attention and stillness for longer. Moreover, dogs rarely manage to perform more than two such runs in one scanner session. These three constraints limit the amount of data that can be collected within a reasonable time-frame. This increases the demands on the data, stressing the importance of data quality. In this report, we describe a hardware approach to circumvent data quantity limitations by increasing data quality.

## 2. Methods

### 2.1. Study Rationale

One avenue to improve data quality is to focus on the “software” side of data analysis, e.g. optimizing data preprocessing, by taking into account the different physiology of dog skulls and brains. Increased data quality was obtained with an inhouse preprocessing pipeline based on SPM, as well as with determining a dog-tailored hemodynamic response function for fMRI analysis (Boch et al., 2021). Another path to improve data quality and analysis sensitivity is the improvement of hardware, through specific dog-tailored hardware components, an avenue that has received less attention thus far.

Dog fMRI usually relies on human scanner systems, which cannot be easily replaced or exchanged to better fit the canine anatomy. Hence, we reasoned that data quality improvements through hardware can be achieved most straightforwardly and cost-effectively through a dog-tailored head coil.

We validated a novel inhouse 16-channel receive coil (K9 coil; distributed by ALSIX GmbH, Austria), which is tailored to the dog’s cranial anatomy (Figure 1). In collaboration with the other co-authors, this coil was developed by CW and EL at the Medical University Vienna. Our intention was to overcome the limitation of commonly used coils (human knee coils, e.g., Jia et al., 2016; Thompkins et al., 2016; Karl et al., 2020, 2021, as well as FlexCoils, e.g. Cuaya et al., 2016; Szabo et al., 2019), which are not tailored to the anatomy of the dog’s skull and thus may result in sub-optimal signal-to-noise ratios and data quality overall.

**Figure 1:**
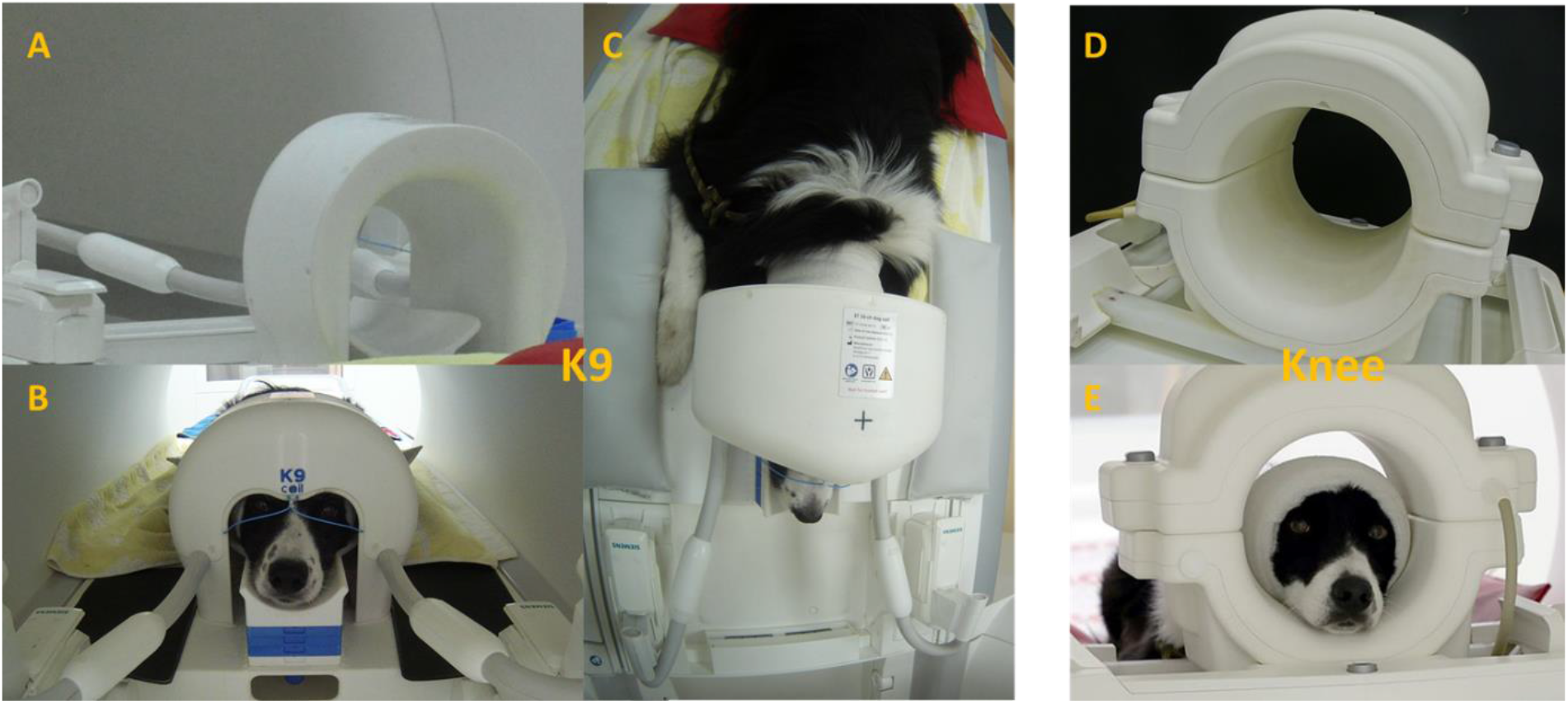
A) rear view of the K9 coil on the scanner bed. B) Front view with subject. Note the chin rest of adaptable height and the paws left and right of the coil. C) Bird’s eye view of dog lying in K9 coil on the scanner bed. D) Rear view of the knee coil. E) Front view of the knee coil with participant. Note the sizable distance between the top of the head and the coil, which is likely reducing sensitivity of measurements.

In the present paper, we apply the K9 coil and compare its images and image quality to a commonly used human knee coil (15 channel receive coil; Siemens Healthineers, Germany) we previously used to scan the same animals (Boch et al., 2021; Karl et al., 2021). To this end, we collected data from nine dogs in three different imaging modalities (structural, functional: task based, functional: resting-state), with the two different coils, using otherwise identical MR scanning parameters.

### 2.2. Sample

Dogs were recruited through the Clever Dog Lab at the Messerli Research Institute at the University of Veterinary Medicine Vienna. Only dogs who completed scanning with both coils were included in this comparison. In total, nine dogs were scanned for T1 imaging and in a functional flickering checkerboard condition with both coils. For resting-state measurements, six of the nine dogs were scanned with both coils and included in our analysis of these resting-state data (Table 1). On average, dogs were 8.1 years old (T1 and functional, 8.3 years in resting-state; note that part of the functional data with the human knee coil and with a different analysis focus was reported already in Boch et al., 2021). Most scanned dogs belonged to herding dog breeds, see Table 1. All dogs had been examined for potential problems with eyesight and general health condition. Dog owners did not receive any monetary compensation for their dogs’ participation and gave written informed consent prior to data collection. All participants in this sample underwent extensive scanner training, based on reward-based positive reinforcement and operant conditioning (Karl et al., 2019), which enabled them to lie unrestrained and still in the MRI scanner. If uncomfortable, dogs are able and allowed to interrupt the run and leave the coil and move on the scanner bed at any time during the examinations, upon which the trainer will give the dog a short break, if so needed, or stop scanning for that day. The studies from which data for this comparative coil overview is taken were approved by the institutional ethics and animal welfare commission in accordance with Good Scientific Practice (GSP) guidelines and national legislation at the University of Veterinary Medicine Vienna (ETK-06/06/2017), based on a pilot study conducted at the University of Vienna. The current study complies with the ARRIVE Guidelines (Kilkenny et al., 2010).

**Table 1:** Demographic data of dogs included in the coil comparison. Note: age indicates age at latest scan. The K9 coil only came into use in 2020, while scanning of the dogs using the knee coil began in 2018. RS = resting-state. All dogs had an 8 minute RS run with each coil, except for Linus who had a 6 minute run.

| name | sex | age | breed | Weight (kg) | T1 | Functional | RS |
| --- | --- | --- | --- | --- | --- | --- | --- |
| Velvet | f | 5 | Labrador Retriever | 26 | x | x | x |
| Maeva | f | 9 | Border Collie/Australian Shepherd Mix | 16 | x | x | x |
| Amy | f | 10 | Border Collie | 23 | x | x | x |
| Emily | f | 12 | Border Collie | 16 | x | x | x |
| Linus | m | 6 | Australian Shepherd | 29 | x | x | x |
| Aeden | m | 12 | Border Collie | 22.5 | x | x | x |
| Cheyenna | f | 6 | Australian Shepherd | 25 | x | x |  |
| Miley | f | 10 | Border Collie | 16 | x | x |  |
| Cameron | m | 7 | Border Collie | 18 | x | x |  |
| **TOTAL** |  | 8.1 |  |  | N = 9 | N = 9 | N = 6 |

### 2.3. Coils

Data and images acquired with the Siemens (human) Tx/Rx 15-channel knee coil were compared to those acquired with the (dog) K9 coil. The K9 imaging coil was designed tailor-made, with special attention to dog head and brain anatomy. The coil is thus composed of 16 linearly polarized receive-only surface channels, 14 of which are mounted inside the coil housing, and two, the “eye-elements”, are partly visible from the outside and consist of flexible cables. The layout of the coil elements, and the flexible rostral elements, were also particularly designed for the larger amount of muscle tissue in the dog’s skull. The coil dimensions were designed with the average size of dogs and dog breeds usually used for neuroimaging in mind, consisting largely of medium sized dogs (mean weight of roughly 20kg), and a high proportion of Border Collies. To bring each dog’s head as close to the inner surface of the coil, an adjustable chin rest was incorporated, allowing for measuring dogs with heads of quite varying sizes (up to 45 cm head circumference), and improving data quality by increasing proximity of the skull to the coil. This tailored chin rest also increases comfort for the dog, making the lying position adaptable to the individual needs of the subject. Additionally, the coil is smaller in width than the human knee coil, allowing the dog to comfortably rest its paws on either side of the coil while its head is inside. An added benefit of higher comfort for the dogs is increased compliance to finish the runs, since dogs will be more reluctant to remain in an uncomfortable setting.

### 2.4. Visual presentation during scanning

For structural imaging (3:12 minutes), dogs were either looking at the trainer sitting in front of the scanner or presented with a video engaging their continuous attention (e.g., showing small animals foraging, such as mice or rooks). The latter approach helped the dogs stay still while they could focus on the screen. During resting-state data acquisition, dogs were presented with a white cross on a black background (run durations between 6 and 8 min, see below). The functional task consisted of 10s blocked presentation of a flickering black and white checkerboard (8Hz) interspersed with 10s cross (green on black background). In total, the run was 2:14 minutes long, including six blocks of visual stimulation and six blocks of baseline in a fixed order, starting with the visual baseline condition.

### 2.5. Data acquisition

Functional imaging data for both the flickering checkerboard task and the resting-state data were obtained from 24 axial slices (interleaved acquisition in descending order, spanning the whole brain) using a 2-fold multiband-accelerated echo planar imaging (EPI) sequence with a voxel size of 1.5 × 1.5 × 2 mm^3^ (TR/TE = 1000/38 ms, field of view (FoV) = 144 × 144 × 58 mm^3^, flip angle = 61°, 20% slice gap). The functional flickering checkerboard task consisted of 134 volumes, the resting-state scans were at least 6 minutes (360 volumes), and at most 8 minutes long (480 volumes), depending on the dog’s capability to lie still for such a prolonged time, without visual input beyond a fixation cross. The structural image was obtained using a voxel size of 0.7 mm isotropic (TR/TE = 2100/3.13 ms, FoV = 230 × 230 × 165 mm^3^). Images in these three modalities were acquired in separate sessions. Note that imaging parameters were chosen to be identical for both coils, so that possible differences in image quality could not be attributed to differences in imaging parameters.

### 2.6. Preprocessing

Preprocessing was run in MATLAB version 2020a, using the SPM12 toolbox. Images were slice-time corrected to the middle slice (see Sladky et al., 2011), and realigned. Thereafter, we performed manual reorientation for the structural and EPI images, and proceeded to manually skull-strip the images with itk-SNAP (Yushkevich et al., 2006). This step is of particular importance in dog MRI, where the skull is bordered by massive musculature which can hinder successful coregistration, which was performed onto the mean image of each run. Structural segmentation of the brain was performed using the canine tissue probability maps provided by (Nitzsche et al., 2019). Normalization of functional and structural data was performed using the “Old Normalization” module in SPM (originally implemented in SPM8), finally reslicing images to 1.5 mm isotropic voxel size, and smoothing of 3 mm (with a Gaussian FWHM kernel). Data were motion scrubbed by calculating framewise displacement, and excluding volumes with a displacement larger than 0.5 mm in comparison to the previous volume (Power et al., 2012, 2014). Roughly 16 volumes had to be excluded on average in the K9 coil, roughly 5 volumes in the knee coil (based on flickering checkerboard runs).

### 2.7. Data analysis

#### 2.7.1. Signal-to-Noise-Ratio (SNR) for structural data

SNR is an important measure of data quality, as it describes the relative contribution of signal of interest vs. noise (of no interest) to the overall recorded signal. One major aim of the K9 coil was to improve SNR by improving signal intensity, foremost by reducing distance between the dog’s brain and the coil elements. We calculated SNR for structural images and temporal SNR for functional images (visual flickering checkerboard and resting-state) using the “SPMUP” toolbox (Pernet, 2014, 2021). This toolbox defines SNR as the ratio between mean signal intensity in the tissue (gray and white matter) by the signal variance outside of the brain, expressed through the standard deviation, or

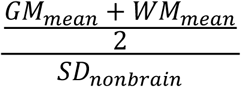

while the tSNR is calculated identically but using the signal over time. For the calculation of SNR, we used unsmoothed and unwarped data. T-Tests and percent differences between coils were calculated using R (version 4.1.0).

#### 2.6.2. Functional fMRI data (resting-state and visual stimulation)

Resting-state data were used to calculate subject-specific tSNR maps. Task data were used to estimate the subject-specific BOLD response to visual stimulation using SPM12’s default settings for a first level single subject t-test (task>0). However, instead of SPM12’s canonical hemodynamic response function (HRF), we used a tailored dog HRF in the analysis of the data (Boch et al., 2021) to account for the faster BOLD response in dogs. The resulting single-subject statistical parametric maps of t-values were transformed into z-values to allow for second-level group analysis. On the group level, we compared tSNR and activation maps between the two coils statistically using paired t-tests in SPM with a threshold of p < 0.05. We used a canine brain atlas (Nitzsche et al., 2019) for parcellation to investigate brain area specific differences.

### 2.8. Data and code availability statement

Data and code can be made available upon written reasonable request to the corresponding authors. The SPMup (https://github.com/CPernet/spmup) and SPM12 toolboxes (https://www.fil.ion.ucl.ac.uk/spm/software/spm12/) are available to the community.

## 3. Results

### 3.1. T1 Data Quality/SNR

For structural images (N = 9), overall group-averaged SNR was 45.28 a.u. (13.78 SD) for the K9 coil, and 31.66 a.u. (15.22 SD) for the knee coil, corresponding to a 43.01% increase of SNR in the K9 coil compared to the knee coil (see Figure 2). The difference was significant, with a large effect size (Cohen’s D = 0.94) and T(8) = 3.98, p < 0.01). We also analyzed SNR for grey and white matter separately. For grey matter, SNR increased (k9 > knee) by 47.03% (T(8) = 4.3, p < 0.005), while it increased by 39.44% for white matter (T(8) = 3.68, p < 0.01) (Cohen’s D = 1.02 and 0.87 respectively).

**Figure 2:**
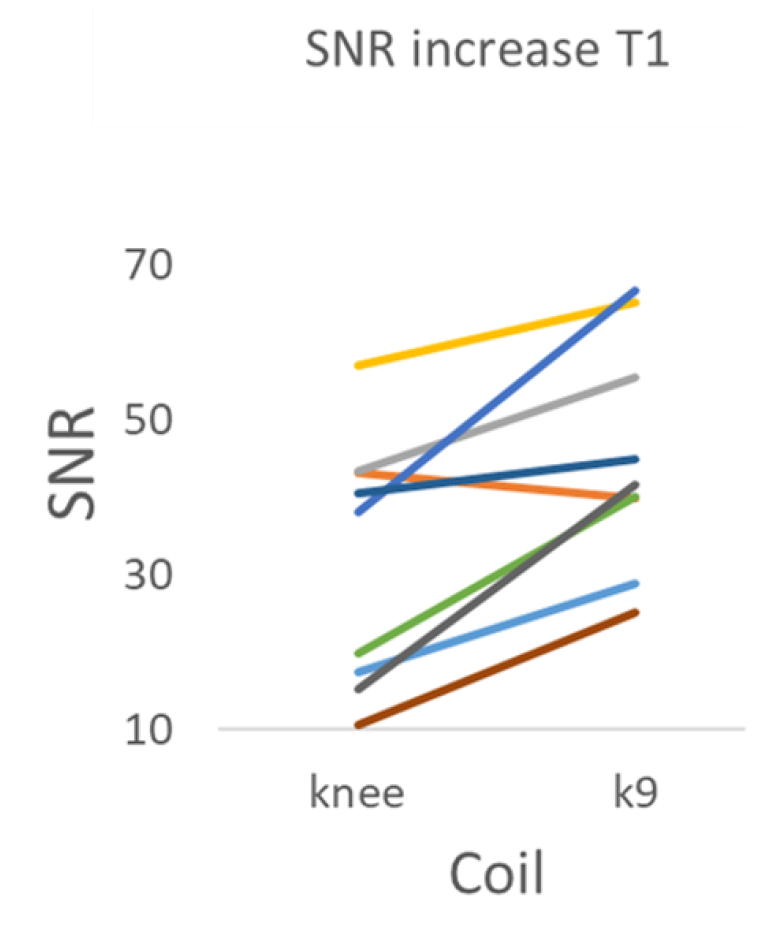
SNR values (AU) for each dog (indicated by different colors) in each coil for the structural data.

### 3.2 Functional neuroimaging: tSNR in resting-state data

We calculated tSNR maps for the K9 and knee coil resting-state data collected in 6 dogs (Figure 3). The K9 coil shows statistically significant tSNR increases in all dorsal brain regions and most ventral brain areas (p < 0.05). No statistically significant tSNR decreases were found. Importantly, no voxels in the knee coil dataset had an increased tSNR when tested with a paired t-test with a threshold of p < 0.05.

**Figure 3:**
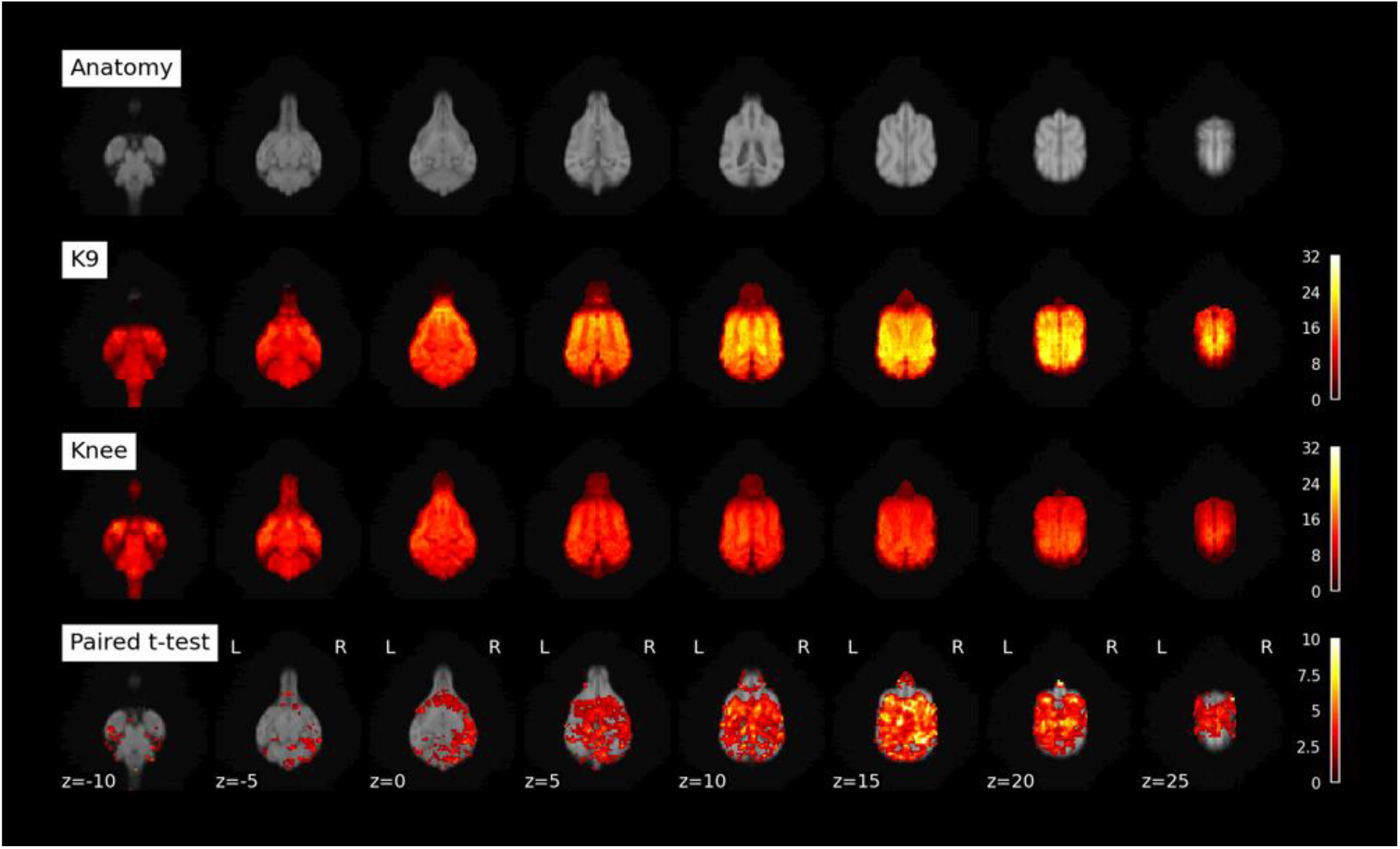
Upper row: anatomical scan from K9 coil T1 images of 6 dogs included in resting-state analysis. K9: tSNR maps for resting-state data collected with K9 coil. Knee: tSNR maps for resting-state data collected with knee coil (both unsmoothed data). Paired t-test: contrasting K9 > Knee (smoothed data).

To quantify the region-specific tSNR increases we performed a comparison based on mean values from a brain parcellation (Nitzsche et al., 2019). In line with the voxel-based analysis, the overwhelming majority of atlas areas showed a statistically significant increase, while no statistically significant tSNR decreases were found (paired t-test, p < 0.05 one-sided; Figure 4 and Table 2). Importantly, over the whole cortex (see encephalon, Table 2) there was a 46.5% increase in tSNR from knee to K9 coil. Some minor decreases were noted in the olfactory bulb, among a few other regions (see Table 2, negative t-values and discussion).

**Table 2:** tSNR differences between K9 and knee coil based on mean regional tSNR for brain parcellations derived from the Nitzsche canine brain atlas (2019). P-values are uncorrected for multiple comparisons and should be regarded as descriptive.

| id | Label | K9 | Knee | Difference | Paired t-test |
| --- | --- | --- | --- | --- | --- |
| 1 | encephalon | 71.12±9.25 | 48.54±8.37 | 46.50% | t=4.0, p=0.005 ** |
| 2 | gyrus frontalis L | 21.90±4.51 | 19.69±2.24 | 11.20% | t=1.1, n.s. |
| 3 | gyrus frontalis R | 23.59±4.20 | 20.92±2.64 | 12.80% | t=1.5, n.s. |
| 4 | gyrus proreus L | 39.34±6.95 | 31.16±5.50 | 26.30% | t=2.1, p=0.043 * |
| 5 | gyrus proreus R | 40.70±7.21 | 30.67±6.06 | 32.70% | t=2.7, p=0.020 * |
| 6 | gyrus compositus rostralis L | 54.91±8.56 | 38.10±8.90 | 44.10% | t=3.2,<br>p=0.012 * |
| 7 | gyrus compositus rostralis R | 50.06±9.65 | 33.84±9.44 | 48.00% | t=2.7,<br>p=0.022 * |
| 8 | gyrus precruciatu s L | 55.78±17.84 | 33.67±5.60 | 65.70% | t=3.2,<br>p=0.012 * |
| 9 | gyrus precruciatu s R | 55.03±16.83 | 34.50±6.00 | 59.50% | t=3.8,<br>p=0.006 ** |
| 10 | gyrus postcruciatu s L | 69.47±19.81 | 40.46±8.30 | 71.70% | t=4.1,<br>p=0.005 ** |
| 11 | gyrus postcruciatu s R | 67.02±19.83 | 38.30±8.60 | 75.00% | t=4.6,<br>p=0.003 ** |
| 12 | gyrus marginalis L | 52.68±12.11 | 32.99±8.33 | 59.70% | t=3.3,<br>p=0.010 * |
| 13 | gyrus marginalis R | 57.79±12.95 | 37.08±9.02 | 55.80% | t=3.6,<br>p=0.007 ** |
| 14 | gyrus ectomarginalis L | 55.83±8.88 | 36.48±5.84 | 53.10% | t=4.1,<br>p=0.004 ** |
| 15 | gyrus ectomarginalis R | 55.33±8.93 | 37.66±7.40 | 46.90% | t=4.3,<br>p=0.004 ** |
| 16 | gyrus occipitalis L | 38.46±6.53 | 28.81±5.29 | 33.50% | t=2.7,<br>p=0.022 * |
| 17 | gyrus occipitalis R | 40.15±4.62 | 28.53±4.35 | 40.70% | t=3.5,<br>p=0.009 ** |
| 18 | gyrus suprasylvius rostralis L | 73.48±14.66 | 45.25±8.05 | 62.40% | t=5.6,<br>p=0.001 ** |
| 19 | gyrus suprasylvius rostralis R | 70.57±13.12 | 41.48±8.02 | 70.10% | t=6.1,<br>p<0.001 *** |
| 20 | gyrus suprasylvius medius L | 79.57±12.77 | 48.72±6.19 | 63.30% | t=5.6,<br>p=0.001 ** |
| 21 | gyrus suprasylvius medius R | 72.25±10.54 | 44.68±6.96 | 61.70% | t=6.6,<br>p<0.001 *** |
| 22 | gyrus suprasylvius caudalis L | 51.63±12.10 | 42.88±5.37 | 20.40% | t=1.8, n.s. |
| 23 | gyrus suprasylvius caudalis R | 51.21±5.75 | 38.21±8.85 | 34.00% | t=3.6,<br>p=0.008 ** |
| 24 | gyrus ectosylvius rostralis L | 76.31±11.66 | 51.06±6.70 | 49.50% | t=5.2,<br>p=0.002 ** |
| 25 | gyrus ectosylvius rostralis R | 71.89±9.96 | 41.40±10.64 | 73.70% | t=6.7,<br>p<0.001 *** |
| 26 | gyrus ectosylvius medius L | 74.88±11.96 | 48.89±6.35 | 53.20% | t=4.0,<br>p=0.005 ** |
| 27 | gyrus ectosylvius medius R | 67.25±9.21 | 41.17±10.48 | 63.40% | t=6.3,<br>p<0.001 *** |
| 28 | gyrus ectosylvius caudalis L | 58.50±9.36 | 46.21±5.33 | 26.60% | t=2.8,<br>p=0.020 * |
| 29 | gyrus ectosylvius caudalis R | 56.72±7.55 | 41.36±10.41 | 37.20% | t=3.9,<br>p=0.006 ** |
| 30 | gyrus sylvius rostralis L | 63.97±12.48 | 47.63±8.96 | 34.30% | t=2.8,<br>p=0.019 * |
| 31 | gyrus sylvius rostralis R | 64.30±9.62 | 43.36±13.05 | 48.30% | t=2.9,<br>p=0.016 * |
| 32 | gyrus sylvius caudalis L | 59.92±9.77 | 49.91±7.59 | 20.10% | t=1.9, n.s. |
| 33 | gyrus sylvius caudalis R | 60.01±8.71 | 41.84±11.10 | 43.40% | t=3.1,<br>p=0.013 * |
| 34 | gyrus compositus caudalis L | 38.16±6.48 | 37.23±6.71 | 2.50% | t=0.3, n.s. |
| 35 | gyrus compositus caudalis R | 38.55±4.61 | 33.68±4.17 | 14.50% | t=2.1,<br>p=0.046 * |
| 36 | gyrus rectus L | 44.63±11.57 | 37.62±10.46 | 18.60% | t=1.8, n.s. |
| 37 | gyrus rectus R | 40.40±12.39 | 34.89±7.38 | 15.80% | t=1.3, n.s. |
| 38 | gyrus genualis L | 27.72±5.41 | 26.22±5.31 | 5.70% | t=0.6, n.s. |
| 39 | gyrus genualis R | 32.03±8.38 | 28.40±6.68 | 12.80% | t=1.3, n.s. |
| 40 | area subcallosa L | 73.57±14.35 | 49.88±10.36 | 47.50% | t=2.7,<br>p=0.022 * |
| 41 | area subcallosa R | 76.44±13.69 | 53.11±11.18 | 43.90% | t=2.9,<br>p=0.018 * |
| 42 | gyrus cinguli L | 71.84±9.74 | 49.00±5.78 | 46.60% | t=4.9,<br>p=0.002 ** |
| 43 | gyrus cinguli R | 71.17±10.03 | 47.37±6.24 | 50.30% | t=5.7,<br>p=0.001 ** |
| 44 | gyrus presplenialis L | 85.93±16.18 | 54.99±6.71 | 56.30% | t=4.4,<br>p=0.004 ** |
| 45 | gyrus presplenialis R | 88.57±18.25 | 55.16±8.18 | 60.60% | t=5.3,<br>p=0.002 ** |
| 46 | gyrus splenialis L | 67.80±8.24 | 49.41±4.64 | 37.20% | t=3.9,<br>p=0.005 ** |
| 47 | gyrus splenialis R | 65.95±6.90 | 48.93±7.33 | 34.80% | t=4.1,<br>p=0.005 ** |
| 48 | gyrus parahippocampalis L | 56.96±7.63 | 47.19±6.76 | 20.70% | t=2.4,<br>p=0.029 * |
| 49 | gyrus parahippocampalis R | 56.60±6.48 | 44.56±9.50 | 27.00% | t=2.7,<br>p=0.020 * |
| 50 | hippocampus L | 63.81±8.23 | 51.85±7.54 | 23.10% | t=2.8,<br>p=0.020 * |
| 51 | hippocampus R | 62.45±7.59 | 48.13±11.91 | 29.70% | t=2.7,<br>p=0.022 * |
| 52 | lobus piriformis L | 39.59±5.47 | 37.13±9.25 | 6.60% | t=0.6, n.s. |
| 53 | lobus piriformis R | 43.34±7.44 | 37.81±7.50 | 14.60% | t=1.4, n.s. |
| 54 | tuberculum olfactorium L | 53.39±14.54 | 38.48±12.15 | 38.80% | t=2.4,<br>p=0.032 * |
| 55 | tuberculum olfactorium R | 52.54±14.80 | 37.57±10.68 | 39.90% | t=2.3,<br>p=0.034 * |
| 56 | gyrus diagonalis L | 56.42±11.43 | 45.23±12.86 | 24.70% | t=1.5, n.s. |
| 57 | gyrus diagonalis R | 60.58±11.06 | 46.82±12.74 | 29.40% | t=1.9, n.s. |
| 58 | gyrus paraterminalis L | 74.58±12.52 | 53.10±12.88 | 40.40% | t=2.2,<br>p=0.041 * |
| 59 | gyrus paraterminalis R | 74.55±12.98 | 56.88±13.81 | 31.10% | t=1.9, n.s. |
| 60 | gyrus olfactorius lateralis L | 52.17±13.00 | 39.86±12.57 | 30.90% | t=1.8, n.s. |
| 61 | gyrus olfactorius lateralis R | 51.04±13.72 | 36.55±11.01 | 39.70% | t=2.4,<br>p=0.032 * |
| 62 | thalamus L | 66.75±8.05 | 53.81±11.14 | 24.00% | t=1.8, n.s. |
| 63 | thalamus R | 66.59±8.08 | 52.51±12.69 | 26.80% | t=1.9, n.s. |
| 64 | bulbus olfactorius L | 10.99±5.86 | 17.01±6.94 | -35.40% | t=-1.7, n.s. |
| 65 | bulbus olfactorius R | 10.57±6.05 | 17.66±6.55 | -40.20% | t=-2.1, n.s. |
| 66 | nucleus caudatus L | 75.36±10.79 | 51.47±10.61 | 46.40% | t=3.5,<br>p=0.009 ** |
| 67 | nucleus caudatus R | 75.36±9.77 | 48.97±11.63 | 53.90% | t=3.6,<br>p=0.008 ** |
| 68 | insular cortex L | 68.19±12.41 | 55.40±9.17 | 23.10% | t=1.7, n.s. |
| 69 | insular cortex R | 70.23±12.95 | 51.89±15.36 | 35.30% | t=2.1,<br>p=0.044 * |
| 70 | hypophysis | 23.82±7.45 | 22.60±6.68 | 5.40% | t=0.3, n.s. |
| 71 | vermis cerebelli | 44.52±7.29 | 36.69±5.93 | 21.30% | t=2.3,<br>p=0.033 * |
| 72 | pons | 30.53±2.94 | 35.31±3.63 | -13.50% | t=-2.4, n.s. |
| 73 | medulla oblongata | 27.76±4.16 | 32.83±5.58 | -15.40% | t=-2.1, n.s. |
| 74 | medulla spinalis | 21.87±4.04 | 26.61±5.57 | -17.80% | t=-1.8, n.s. |
| 75 | mesencephalon | 45.48±5.12 | 44.95±6.65 | 1.20% | t=0.1, n.s. |
| 76 | diencephalon | 55.70±7.70 | 48.68±9.75 | 14.40% | t=1.2, n.s. |
| 77 | nervus opticus | 38.74±12.28 | 31.93±8.79 | 21.30% | t=1.2, n.s. |
| 78 | hemispherium cerebelli L | 35.54±6.17 | 32.37±5.65 | 9.80% | t=1.3, n.s. |
| 79 | hemispherium cerebelli R | 36.75±4.89 | 31.20±4.28 | 17.80% | t=3.1,<br>p=0.013 * |
| 80 | commissura rostralis | 71.08±10.63 | 58.07±14.46 | 22.40% | t=1.4, n.s. |
| 81 | pedunculus olfactorius L | 31.04±8.41 | 30.13±14.68 | 3.00% | t=0.2, n.s. |
| 82 | pedunculus olfactorius R | 31.34±11.36 | 30.75±10.45 | 1.90% | t=0.2, n.s. |
| 83 | area septalis L | 77.09±13.20 | 49.41±10.64 | 56.00% | t=3.5,<br>p=0.009 ** |
| 84 | area septalis R | 77.54±13.71 | 51.78±12.53 | 49.70% | t=3.2,<br>p=0.012 * |
| 85 | nucleus et tractus spinalis nervi trigemini L | 27.28±3.49 | 32.52±7.18 | -16.10% | t=-2.1, n.s. |
| 86 | nucleus et tractus spinalis nervi trigemini R | 28.66±5.24 | 32.53±6.39 | -11.90% | t=-1.4, n.s. |
| 87 | nucleus ventralis caudalis thalami pars medialis L | 61.75±9.00 | 58.64±13.84 | 5.30% | t=0.4, n.s. |
| 88 | nucleus ventralis caudalis thalami pars medialis R | 59.68±9.03 | 55.57±14.06 | 7.40% | t=0.5, n.s. |
| 89 | amygdala L | 47.08±7.24 | 49.34±11.89 | -4.60% | t=-0.4, n.s. |
| 90 | amygdala R | 48.58±8.94 | 46.20±9.44 | 5.10% | t=0.5, n.s. |

**Figure 4:**
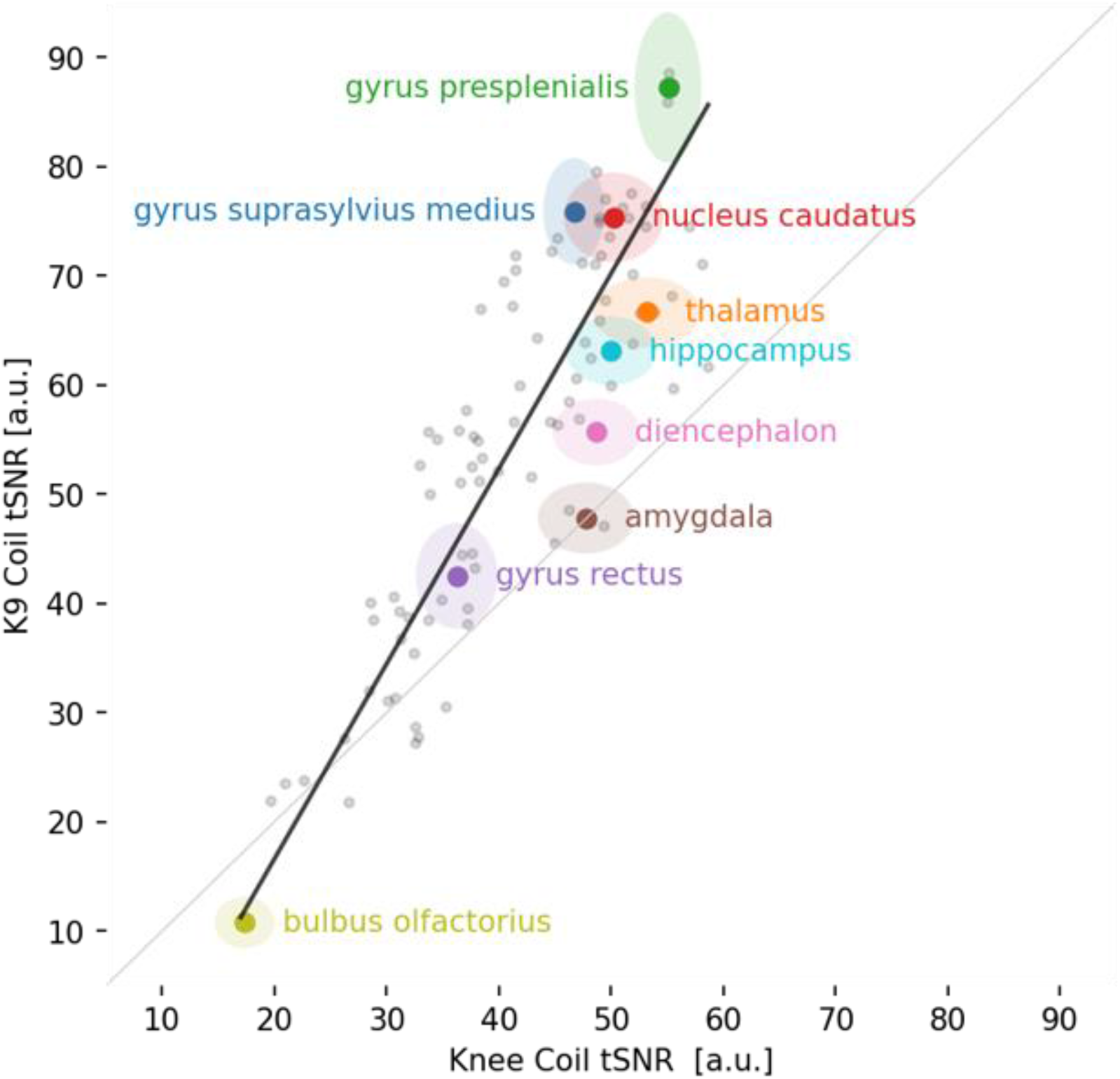
Scatterplot of 90 brain areas averaged across all 6 dogs in the analysis. Almost all brain areas (visualized as grey dots) fall above the grey identity line, hence showing a tSNR increase for the K9 coil. Some important brain areas of interest are color-coded, labeled, and displayed with their 95% confidence intervals.

Figure 4 gives an overview of all brain areas and their change in tSNR from the Knee to the K9 coil in the resting-state data.

### 3.3 Functional neuroimaging: Activation in the visual flickering checkerboard

For the visual flickering checkerboard, we had data from both coils from 9 dogs. Contrasting activation to baseline, we found activation in the visual cortex with both coils (see Figure 5, top two rows). A paired t-Test (bottom row, K9 > Knee) shows areas in which the K9 coil outperformed the knee coil in the visual cortex.

**Figure 5:**
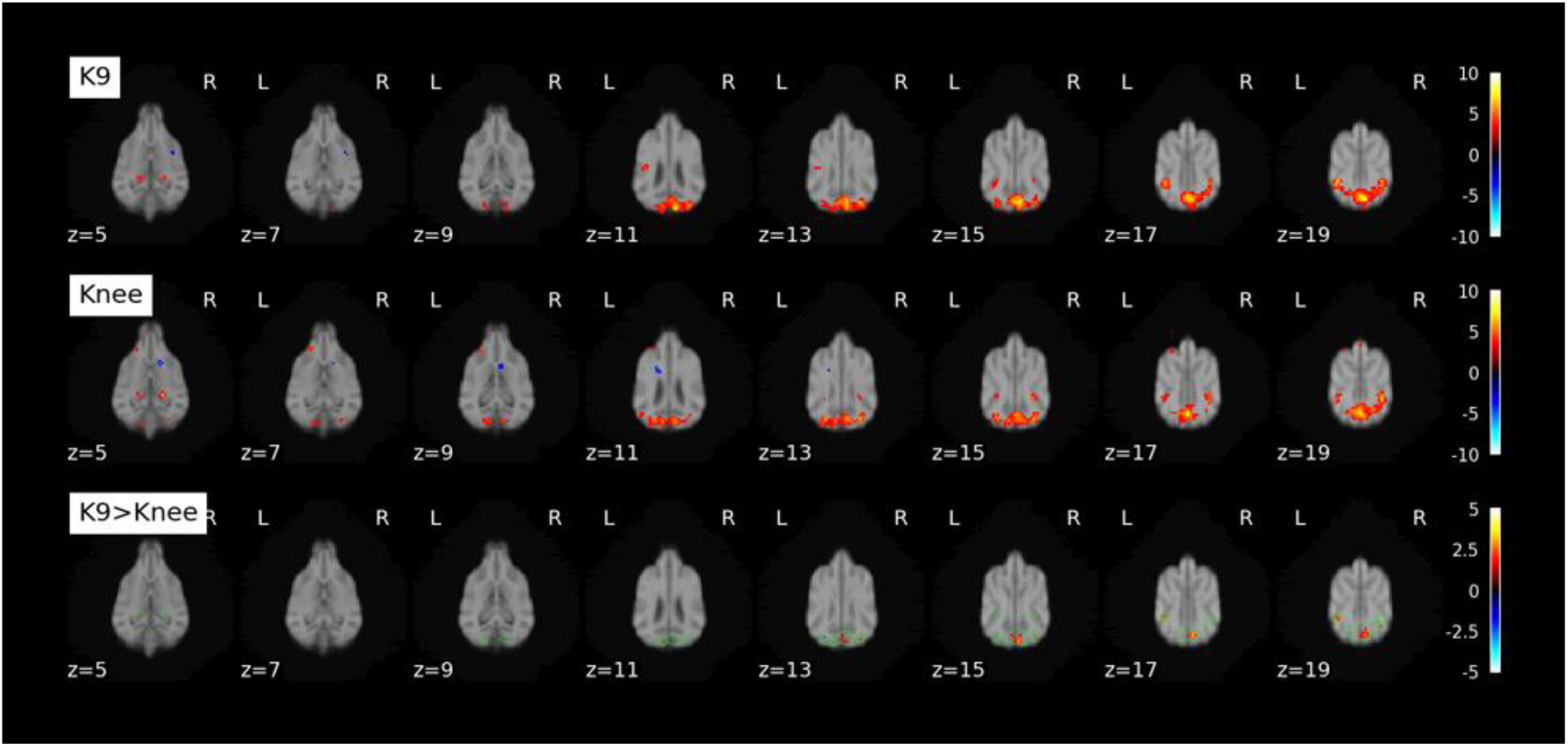
K9, top row: Activation found contrasting Checkerboard > Baseline (fixation cross) with the K9 coil. Knee, middle row: Activation found contrasting Checkerboard > Baseline (fixation cross) with the knee coil. K9 > Knee, bottom row: paired t-Test. Green outlines show conjunction of K9 and Knee coil activation. Second-level analysis was performed on the single-subject contrasts and thresholded at p<0.001 (k>=5 voxels for display purposes).

Furthermore, we looked at individual changes in z-scores in each voxel in all the 9 dogs included in the analysis of the visual flickering checkerboard. Most, but not all, dogs’ signal improved with the K9 coil (see Figure 6), and increases in z-scores were mostly larger than decreases.

**Figure 6:**
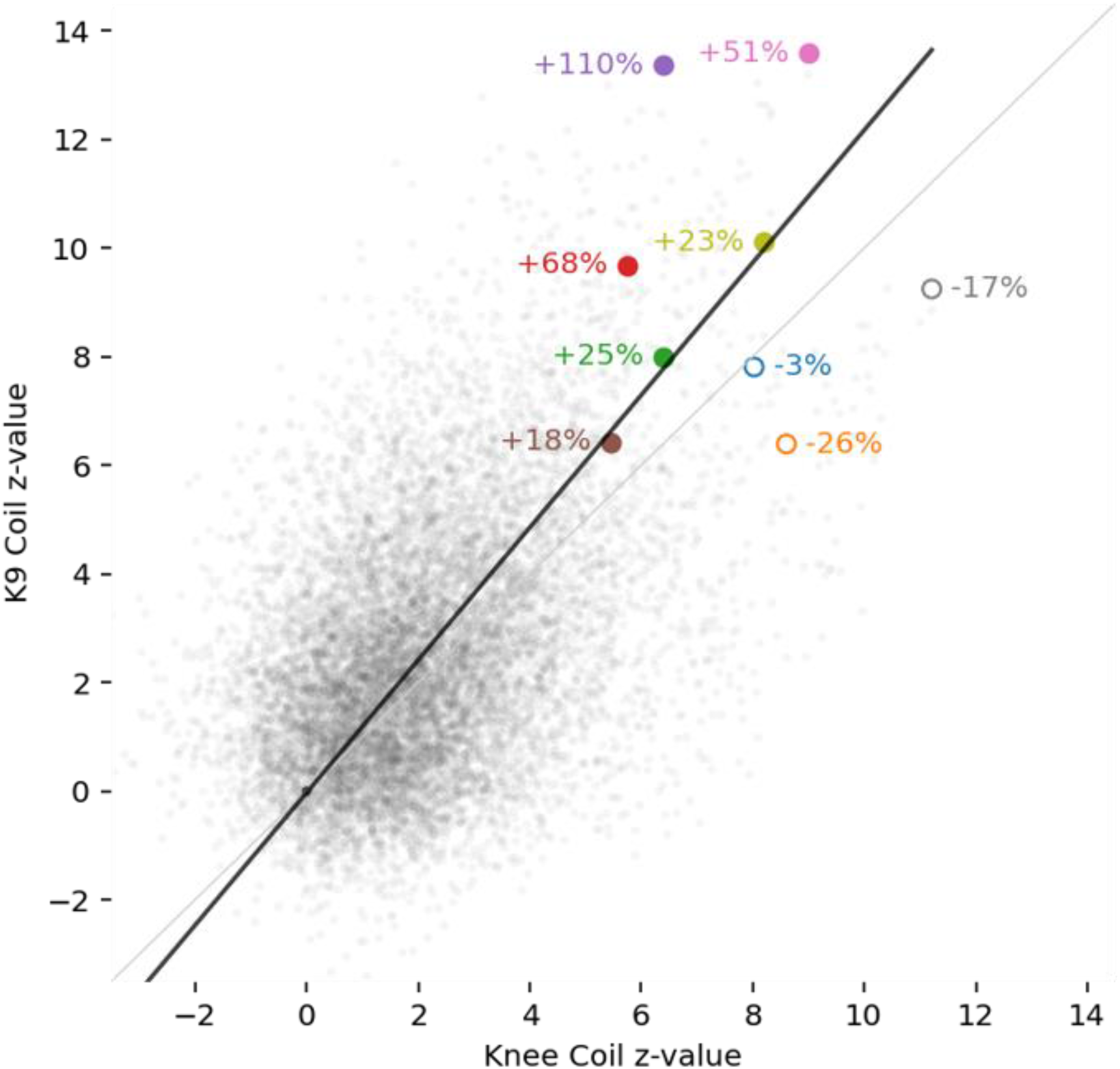
Individual z-values for each voxel in each dog in the data from the visual flickering checkerboard paradigm. Largest individual increases (6 dogs) or decreases (3 dogs) are labeled.

### 3.4 Movement correlation

Degree of correlation between movement and signal did not differ systematically between the coils (all p > 0.05), and neither did raw framewise displacement (all p > 0.2).

## 4. Discussion

The aim of this study was to validate the new K9 coil across various MRI modalities. To this end, we compared results from the K9 coil with results from a human knee coil, commonly used for dog fMRI. Data were compared in terms of data quality as expressed in SNR, and second level results in a classical GLM fMRI analysis. Since the design of the K9 coil was tailored to dog cranial anatomy, we expected the K9 coil data quality to outperform the knee coil, and possibly lead to more robust results.

The comparison of the standard human knee coil for dog brain imaging with our inhouse K9 coil has produced a range of evidence that the K9 coil indeed offers higher sensitivity compared to the knee coil. In particular, spatial and temporal signal-to-noise ratios were increased with the K9 coil, across all imaging modalities. In the structural data, we noted an increase of roughly 45% across grey and white matter. Of note, since the K9 coil came into use later, dogs might have been more trained but also older. The expected increase in SNR (and tSNR) due to better training should however be more than mitigated by increases in signal noise that are observable with increased age (in humans, McIntosh et al., 2013; Yao et al., 2013).

In functional imaging modalities, the differences were also very noticeable. With regard to the resting-state scans, both the knee and the K9 coil can be used for dog fMRI, however the K9 coil was much more sensitive in terms of both SNR and tSNR across the canine brain in our small sample of dogs (n = 6 for resting-state). All dorsal brain areas exhibited increases in tSNR in the K9 coil, and this is of particular interest for the investigation of convergent evolution of higher cognition, such as social cognition, since dorsal areas are more likely to contribute to these operations (Rushworth et al., 2013). While Figure 4 shows also decreases of tSNR in the K9 coil as compared to the knee coil, most notably in the olfactory lobe, no voxel was found to have statistically better tSNR in the knee coil as compared to the K9 coil. Finally, tSNR in the resting-state modality saw a similar increase as in the structural modality, of roughly 46% from knee to K9 coil. Please note that some of the decreases noted could also stem from changes in the field of view settings: we had issues with wrapovers in the temporal lobe, and fixed this by tilting the field of view, so that in some dogs, parts of the olfactory bulb might have been cut.

For the visual flickering checkerboard, we found robust activation in the primary visual cortex of dogs with both coils. However, with the K9 coil, a few additional clusters were identified in the paired t-test, in particular in the occipital lobe, as can be seen in Figure 5. On an individual level, not all dogs benefitted from the K9 coil equally, some even had decreases in voxelwise z scores (Figure 6). However, fewer individuals exhibited decreases, and the decreases were generally smaller than the increases found in the other dogs. The tSNR increases seen across modalities thus also translated into more activation being detected in highly plausible areas (occipital lobe, primary visual cortex) in a standard second level GLM analysis of fMRI data in a robust paradigm.

Overall, the strongest evidence in favor of the K9 coil comes from the raw SNR and tSNR increases. These clearly demonstrate that data quality is much improved in the K9 coil. Based on the lack of differences in raw framewise displacement between the coils, this difference does not solely come from a reduction in motion artifacts, but rather directly from the coil properties. The substantial improvements in SNR do also lead to improvements on the second level analysis of the functional visual flickering checkerboard data. Despite the visual flickering checkerboard paradigm being very robust, we were still able to find a multitude of small clusters of brain areas that were only present with the K9 coil. The increase in tSNR in all dorsal and most ventral regions with the K9 coil, our new hardware offers the opportunity to investigate smaller effects of interest, which is particularly relevant for the investigation of higher order cognition, as well as social cognition, in dogs and potentially other canines.

The main aim of this study was to examine possible benefits of a dog-tailored MR coil. We find compelling evidence that the K9 coil will lead to significant improvements in data quality and dog MR imaging. It should be noted though that the K9 coil comes with its own shortcomings due to its high specificity: it is limited to usage in dogs, not all canines, and tailored to a reduced range of breeds in particular. Some larger-skulled dogs will not fit, and for very small dogs the distance to the coil elements might also be too great. But the same would hold true and even more so for the human knee coil.

The K9 coil yields an almost 50% increase in SNR compared to the knee coil, in particular in dorsal cortical areas, across all investigated modalities. With canine neuroimaging as an emerging field, key constraints of small samples and short functional runs emphasize the need for tailored hardware. While existing human imaging hardware will certainly lend valid results as well, especially when robust effects can be expected, the K9 coil offers improved data quality, better subject fit and comfort, and we thus expect it to be a key contribution to the ongoing advancement of dog and canine neuroimaging.

## Acknowledgements

This project was funded in part by the Austrian Science Fund (FWF) [P33180] and by the Vienna Science and Technology Fund (WWTF), the City of Vienna and ithuba Capital AG through project CS18-012, and the Messerli Foundation (Sörenberg, Switzerland). The funders had no role in study design, data collection and analysis, decision to publish, or preparation of the manuscript.

## References

Behroozi M (2019) Establishing a novel fMRI approach to investigate visual cognitive properties in pigeons.

Behroozi M, Billings BK, Helluy X, Manger PR, Güntürkün O, & Ströckens F (2018) Functional MRI in the Nile crocodile: A new avenue for evolutionary neurobiology. Proceedings of the Royal Society B: Biological Sciences, 285(1877), 20180178.

Behroozi M, Helluy X, Ströckens F, Gao M, Pusch R, Tabrik S, Tegenthoff M, Otto T, Axmacher, N, Kumsta R, Moser D, Genc E, & Güntürkün O (2020) Event-related functional MRI of awake behaving pigeons at 7T. Nature Communications, 11(1), 4715. https://doi.org/10.1038/s41467-020-18437-1

Berns G (2013) How dogs love us: A neuroscientist and his adopted dog decode the canine brain. Houghton Mifflin Harcourt.

Berns GS, Brooks AM, & Spivak M (2012) Functional MRI in Awake Unrestrained Dogs. PLOS ONE, 7(5), e38027. https://doi.org/10.1371/journal.pone.0038027

Boch M, Karl S, Sladky R, Huber L, Lamm C, & Wagner IC (2021) Tailored haemodynamic response function increases detection power of fMRI in awake dogs (Canis familiaris). NeuroImage, 224, 117414. https://doi.org/10.1016/j.neuroimage.2020.117414

Bunford N, Andics A, Kis A, Miklósi Á, & Gácsi M (2017) Canis familiaris As a Model for Non-Invasive Comparative Neuroscience. Trends in Neurosciences, 40(7), 438–452. https://doi.org/10.1016/j.tins.2017.05.003

Cuaya LV, Hernández-Pérez R, & Concha L (2016) Our Faces in the Dog’s Brain: Functional Imaging Reveals Temporal Cortex Activation during Perception of Human Faces. PLOS ONE, 11(3), e0149431. https://doi.org/10.1371/journal.pone.0149431

Güntürkün O & Bugnyar T (2016) Cognition without cortex. Trends in cognitive sciences, 20(4), 291–303.

Huber L & Lamm C (2017) Understanding dog cognition by functional magnetic resonance imaging. Learning & Behavior, 45(2), 101–102. https://doi.org/10.3758/s13420-017-0261-6

Jia H, Pustovyy OM, Wang Y, Waggoner P, Beyers RJ, Schumacher J, Wildey C, Morrison E, Salibi N, Denney TS, Vodyanoy VJ, & Deshpande G (2016) Enhancement of Odor-Induced Activity in the Canine Brain by Zinc Nanoparticles: A Functional MRI Study in Fully Unrestrained Conscious Dogs. Chemical Senses, 41(1), 53–67. https://doi.org/10.1093/chemse/bjv054

Kaminski J & Nitzschner M (2013) Do dogs get the point? A review of dog–human communication ability. Learning and Motivation, 44(4), 294–302.

Karl S, Boch M, Virányi Z, Lamm C, & Huber L (2019) Training pet dogs for eye-tracking and awake fMRI. Behavior research methods, 1–19.

Karl S, Boch M, Zamansky A, van der Linden D, Wagner IC, Völter CJ, Lamm C, & Huber L (2020) Exploring the dog–human relationship by combining fMRI, eye-tracking and behavioural measures. Scientific Reports, 10(1), 22273. https://doi.org/10.1038/s41598-020-79247-5

Karl S, Sladky R, Lamm C, & Huber L (2021) Neural Responses of Pet Dogs Witnessing their caregiver’s Positive Interactions with a Conspecific: An fMRI Study. Cerebral Cortex Communications.

Keilholz SD, Silva AC, Raman M, Merkle H, & Koretsky AP (2004) Functional MRI of the rodent somatosensory pathway using multislice echo planar imaging. Magnetic Resonance in Medicine, 52(1), 89–99. https://doi.org/10.1002/mrm.20114

Kilkenny C, Browne W, Cuthill IC, Emerson M, & Altman DG (2010) Animal research: Reporting in vivo experiments: The ARRIVE guidelines. British Journal of Pharmacology, 160(7), 1577–1579. https://doi.org/10.1111/j.1476-5381.2010.00872.x

Kirchhofer KC, Zimmermann F, Kaminski J, & Tomasello M (2012) Dogs (Canis familiaris), but not chimpanzees (Pan troglodytes), understand imperative pointing. PloS one, 7(2), e30913.

Mars RB, Verhagen L, Gladwin TE, Neubert FX, Sallet J, & Rushworth MF (2016) Comparing brains by matching connectivity profiles. Neuroscience & Biobehavioral Reviews, 60, 90–97.

McIntosh AR, Vakorin V, Kovacevic N, Wang H, Diaconescu A, & Protzner AB (2014) Spatiotemporal Dependency of Age-Related Changes in Brain Signal Variability. Cerebral Cortex, 24(7), 1806–1817. https://doi.org/10.1093/cercor/bht030

Nitzsche B, Boltze J, Ludewig E, Flegel T, Schmidt MJ, Seeger J, Barthel H, Brooks OW, Gounis MJ, Stoffel MH, & Schulze S (2019) A stereotaxic breed-averaged, symmetric T2w canine brain atlas including detailed morphological and volumetrical data sets. NeuroImage, 187, 93–103. https://doi.org/10.1016/j.neuroimage.2018.01.066

Pernet C (2021). SPM UP [MATLAB]. https://github.com/CPernet/spmup (Original work published 2014)

Power JD, Barnes KA, Snyder AZ, Schlaggar BL, & Petersen SE (2012) Spurious but systematic correlations in functional connectivity MRI networks arise from subject motion. Neuroimage, 59(3), 2142–2154.

Power JD, Mitra A, Laumann TO, Snyder AZ, Schlaggar BL, & Petersen SE (2014) Methods to detect, characterize, and remove motion artifact in resting state fMRI. NeuroImage, 84, 320–341. https://doi.org/10.1016/j.neuroimage.2013.08.048

Rilling JK (2014) Comparative primate neuroimaging: Insights into human brain evolution. Trends in cognitive sciences, 18(1), 46–55.

Rushworth MF, Mars RB, & Sallet J (2013) Are there specialized circuits for social cognition and are they unique to humans? Current Opinion in Neurobiology, 23(3), 436–442. https://doi.org/10.1016/j.conb.2012.11.013

Sladky R, Friston KJ, Tröstl J, Cunnington R, Moser E, & Windischberger C (2011) Slice-timing effects and their correction in functional MRI. Neuroimage, 58(2), 588–594.

de Schotten TM, Croxson PL, & Mars RB (2019) Large-scale comparative neuroimaging: Where are we and what do we need? Cortex, 118, 188–202. https://doi.org/10.1016/j.cortex.2018.11.028

Strassberg LR, Waggoner LP, Deshpande G, & Katz JS (2019) Training Dogs for Awake, Unrestrained Functional Magnetic Resonance Imaging. JoVE (Journal of Visualized Experiments), 152, e60192. https://doi.org/10.3791/60192

Szabó D, Czeibert K, Kettinger Á, Gácsi M, Andics A, Miklósi Á, & Kubinyi E (2019) Resting-state fMRI data of awake dogs (Canis familiaris) via group-level independent component analysis reveal multiple, spatially distributed resting-state networks. Scientific Reports, 9(1), 15270. https://doi.org/10.1038/s41598-019-51752-2

Thompkins AM, Deshpande G, Waggoner P, & Katz JS (2016) Functional Magnetic Resonance Imaging of the Domestic Dog: Research, Methodology, and Conceptual Issues. Comparative cognition & behavior reviews, 11, 63–82. https://doi.org/10.3819/ccbr.2016.110004

Xu T, Nenning KH, Schwartz E, Hong SJ, Vogelstein JT, Goulas A, Fair DA, Schroeder CE, Margulies DS, & Smallwood J (2020) Cross-species functional alignment reveals evolutionary hierarchy within the connectome. Neuroimage, 223, 117346.

Yao Y, Lu WL, Xu B, Li CB, Lin CP, Waxman D, & Feng JF (2013) The Increase of the Functional Entropy of the Human Brain with Age. Scientific Reports, 3(1), 2853. https://doi.org/10.1038/srep02853

Yushkevich PA, Piven J, Hazlett HC, Smith RG, Ho S, Gee JC, & Gerig G (2006) User-guided 3D active contour segmentation of anatomical structures: Significantly improved efficiency and reliability. Neuroimage, 31(3), 1116–1128.

